# Subgenomic flaviviral RNAs and human proteins: *in silico* exploration of anti-host defense mechanisms

**DOI:** 10.1101/2024.06.05.597601

**Authors:** Riccardo Delli Ponti, Andrea Vandelli, Gian Gaetano Tartaglia

## Abstract

**Background:** Flaviviruses pose significant global health threats, infecting over 300 million people annually. Among their evasion strategies, the production of subgenomic flaviviral RNAs (sfRNAs) from the 3’ UTR of viral genomes is particularly notable. These sfRNAs interact with human proteins, disrupting key cellular processes such as RNA splicing and the interferon response.

**Results:** Utilizing a comprehensive *in silico* approach with the *cat*RAPID algorithm, we analyzed over 300,000 interactions between sfRNAs and human proteins derived from more than 8,000 flavivirus genomes, including Dengue, Zika, Yellow Fever, West Nile, and Japanese Encephalitis viruses. Our study not only validated known interactions but also revealed novel human proteins that could be involved in sfRNA-mediated host defense evasion, including helicases, splicing factors, and chemokines. We propose that sfRNAs function as molecular sponges, sequestering specific proteins indicative of sfRNA-forming regions across flaviviruses. These findings represent a valuable resource for diagnostic and therapeutic developments.

**Conclusions:** Our findings significantly expand the known interactome of sfRNAs with human proteins, underscoring their role in modulating host cellular pathways. By providing the first extensive atlas of sfRNA interactions, we offer new insights into how flaviviruses can manipulate host cellular machinery to facilitate viral survival and persistence. Intriguingly, we predict interaction with stress granules, a critical component of the cellular response to viral infection, suggesting a mechanism by which flaviviruses inhibit their formation to evade host defenses. This atlas not only serves as a resource for exploring therapeutic targets but also aids in the identification of sfRNA biomarkers for improved flavivirus diagnostics.

## Background

Flaviviruses are a class of single-stranded RNA viruses including Dengue virus (DENV), Zika virus (ZIKV), Yellow Fever virus (YFV), West Nile virus (WNV) and Japanese Encephalitis virus (JEV). The flaviviral genome comprises one open-reading frame, encoding for 10 genes flanked by 5’ and 3’ UTR [1, 2]. Several flaviviruses are arboviruses having mosquitoes as intermediary hosts. While flaviviruses are more highly predominant in tropical environments, global warming is moving the threat toward Europe and North America. DENV has already been detected as endemic in different European countries, while West Nile virus (WNV) has been endemic in the USA since 1999 [3]. According to recent estimates, >300 million people are in danger of potentially contracting Dengue virus, with >100 million infections every year [4–6]. RNA viruses have different mechanisms to disrupt the human cellular machinery and innate immune response to guarantee their fitness. Flaviviruses are not an exception, with different mechanisms to avoid the host-defense systems. DENV and hepatitis C virus (HCV) induce rearrangements inside the cellular membrane to compartmentalize their replication machinery and regulate the access of antiviral host proteins [7, 8]. Moreover, the majority of flaviviruses, including DENV, HCV, and YFV, can disrupt the interferon (IFN) signaling cascade by cleaving or interacting with the STING protein [9–11]. The viral infection also triggers and alters different cellular mechanisms. RNA splicing was shown to be altered after ZIKV infection, which generates alternative splicing events in >200 RNAs [12]. Other cellular mechanisms, including RNA editing and decay, are potentially involved in anti-viral response, thus becoming a target to be disrupted by viruses [13–15].

During infection, flaviviruses not only generate copies of their genomic RNA (gRNA) but also smaller RNAs, the subgenomic flaviviral RNAs (sfRNAs). Compared to the gRNAs (∼11Kb), sfRNAs are much smaller, around 300-500 nucleotides [16, 17]. The sfRNAs are viral fragments generated at the 3’ UTR of the viral genome. The mechanism involves the 5′-3′ exoribonuclease XRN1, a host-specific protein that binds and progressively digests the viral genome. However, flaviviral genomes possess specific complex and rigid stem-loops (SL) in their 3’ UTR that stall XRN1, especially SL-II, thus preventing further digestion [17–19]. These XRN1-resistant structures are also present in the mosquito vector and tend to be conserved in different flaviviruses [19, 20]. The existence of these XRN1-resistant structural elements allows the accumulation of the sfRNAs, partially digested RNA fragments at the 3’ UTR of the flaviviral genomes. The sfRNAs are non-coding RNAs (ncRNAs), however, their existence and cellular presence is correlated with the virulence and pathogeny of each flavivirus [3, 16].

The presence of sfRNAs is essential for the pathogenicity of WNV, since mutants lacking sfRNAs were poorly replicating in mice [3, 19]. Moreover, the intricated secondary structures at the 3’ UTR are essential for the formation and functionality of sfRNAs [19]. Due to their high-concentration and complex secondary structures, it is speculated that sfRNAs can act as protein sponges with an anti host-defence function. Different studies focused on characterizing the human proteins binding to the sfRNA or the 3’ UTR, especially of DENV and ZIKV [21, 22]. The cellular functions disrupted by the binding of human proteins with sfRNAs includes RNAi and innate immunity, especially the interferon response. However, experimental works mainly focused on DENV and ZIKV, usually only using a representative genome of each, with few high-binding protein candidates in common among the different studies.

In this work, we used >8000 flaviviral genomes coming from DENV, ZIKV, WNV, JEV and YFV to generate >300,000 in-silico interactions between sfRNAs and human proteins. Our objective is to study the ability of sfRNAs to interfere with the human RBP network. We were able to identify several mechanisms altered by the binding of human proteins with sfRNAs, categorizing species- and couple-specific proteins between the 5 different flaviviruses. We propose that sfRNAs act as *protein sponges* establishing strong interactions with human RNAs. We identified a core set of 21 proteins in common between all the 5 sfRNAs, mainly involving RNA helicases and their interactors. These proteins exhibit a predictive power to identify sfRNA-forming regions in other flaviviruses, and can be used for further investigations through the *cat*RAPID *omics* algorithm.

## Methods

### Sequence and structural studies

We used *CD-HIT* [23] to reduce the sequence redundancy at 90% for all the sfRNAs in our set. We then used *Emboss needleall* [24] and *Mafft* [25] to respectively compute pairwise sequence identity and build a multiple-alignment. The *CROSS* algorithm [26], with the *Global Score* module, was used to predict secondary structure profiles. The profiles were then employed to extract a secondary structure consensus profile by averaging *CROSS* score for every position.

### sfRNAs dataset creations

We downloaded the complete genomes of five different flaviviruses: Dengue virus (DENV), Zika virus (ZIKV), West Nile virus (WNV), Japanese Encephalitis virus (JEV), and Yellow Fever virus (YFV). The DENV and ZIKV genomes were sourced from the Virus Pathogen Resource (VIPR), now known as the Bacterial and Viral Bioinformatics Resource Center (Bv-Brc), and comprised over 5,000 and 1,000 genomes respectively. The genomes for WNV, JEV, and YFV were obtained from the National Center for Biotechnology Information (NCBI), as detailed in **Table 1**. After downloading the complete genomes, we filtered out genomes with unknown nucleotides (any number of “N” in their genomes). After that, we selected the last 500 nt at the 3’ UTRs of all the >8000 genomes as representatives of the sfRNAs, accordingly to the coordinates of the XRN1-stalling region in DENV and WNV ([16]; Supplementary Figure 1). These 500 nt fragments were then filtered for sequence similarity using CD-HIT (90% redundancy; [23]).

### Predicting protein-RNA interactions

Interactions between the viral sfRNAs sequences and the human RNA-biding proteome (RBPome) were predicted using *cat*RAPID *omics* [27], an algorithm to estimate the binding propensity of protein–RNA pairs by combining secondary structure, hydrogen bonding and van der Waals contributions. The predictions of the viral sequences against ∼1500 human RNA-binding proteins (RBPs) are available at the following links:

### ZIKV

http://crg-webservice.s3.amazonaws.com/submissions/2021-04/351700/output/index.html?unlock=c3e033d661

### JEV

http://crg-webservice.s3.amazonaws.com/submissions/2021-04/352201/output/index.html?unlock=cdaa7858e1

### YFV

http://crg-webservice.s3.amazonaws.com/submissions/2021-04/352204/output/index.html?unlock=b61079c43e

### DENV

http://crg-webservice.s3.amazonaws.com/submissions/2021-04/351702/output/index.html?unlock=57c29a2684

### WNV

http://crg-webservice.s3.amazonaws.com/submissions/2021-04/351706/output/index.html?unlock=4d5a11192f

The output is filtered according to the *Z*-score, which is the interaction propensity normalized by the mean and standard deviation calculated over the reference RBP set (http://s.tartaglialab.com/static_files/shared/faqs.html#4). We then selected a threshold of Z-score > 1.5 to assess the most relevant interactions, a method employed in previous publications [28]. Consequently, proteins with at least one interaction with a Z-score > 1.5 for a specific sfRNA were considered highly interacting with that virus. Proteins having Z-score > 1.5 only for a selected virus, or the interaction of two viruses, were considered as species- or couple-specific.

### Selecting and comparing experimental proteins

To validate the quality of our predictions, we used a set of known experimental-validated proteins interacting with DENV sfRNA and 3’ UTR [21, 22]. For sfRNA-specific proteins, from the original paper, we selected only the proteins specific for DENV (21 proteins). Regarding the 3’-specific proteins, we selected only the proteins with an enrichment >1.5 (experimental vs control ratio) in any replicate for any DENV serotype, as suggested by the authors of the manuscript (27 proteins). When comparing the interactions between the sfRNAs and the human RBPome, we integrated experimentally validated proteins not present in the original RBPome as custom libraries inside *catRAPID omics* [27]. To check how well these proteins are predicted, we ranked the Z-score of all the predicted interactions (>300,000 interactions between sfRNAs and human proteins). Then, we selected the best 10 interactions for each experimentally validated protein, and we checked how well they ranked in the top ranked percentage of the overall distribution (**Figure 1**).

**Figure 1:**
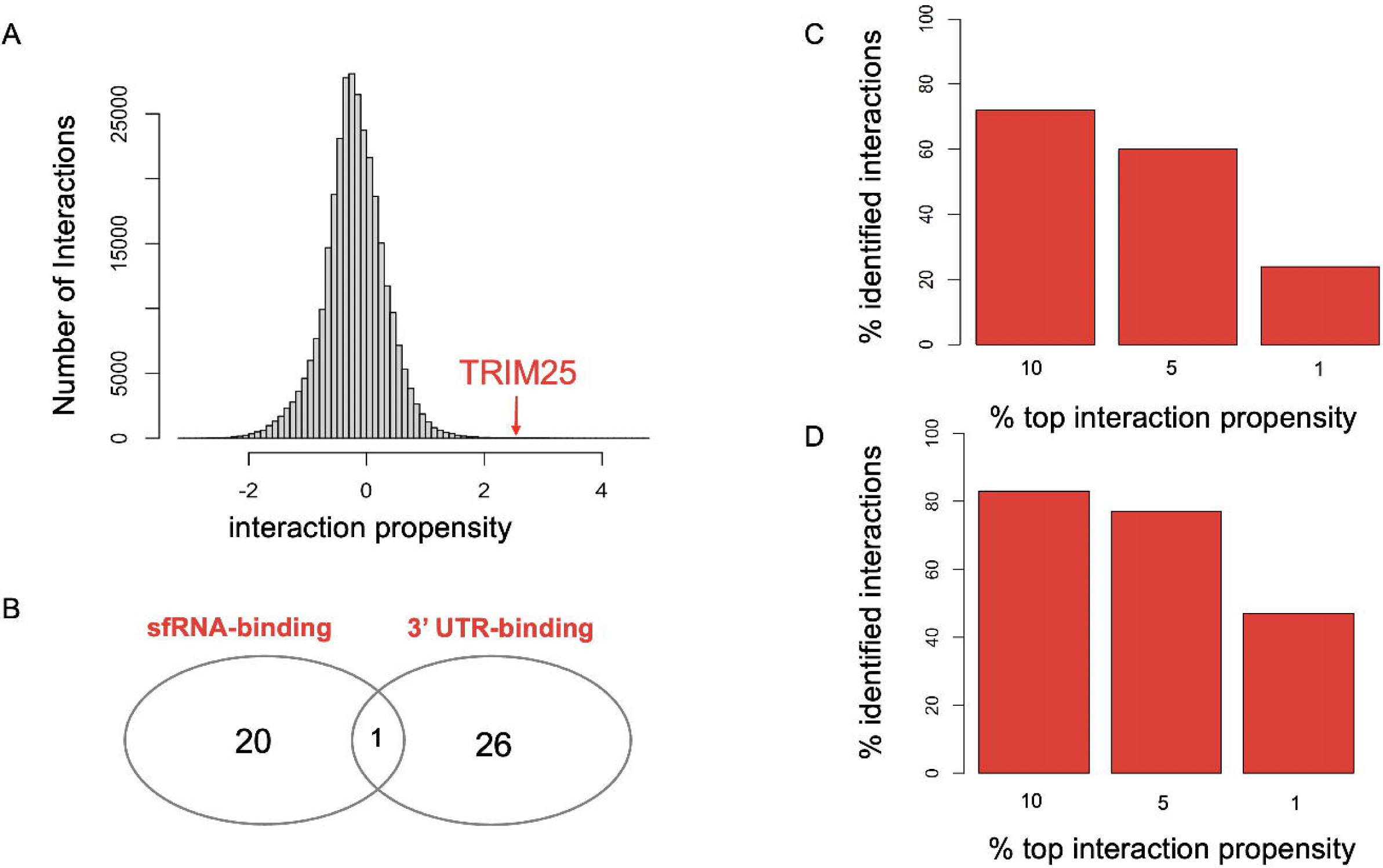
**(A)** Histogram showing the interacting propensity (catRAPID Z-score) between the human proteome and DENV sfRNAs. Interactions with a Z-score > 1.5 are considered high interactions. TRIM25, known interactor of DENV sfRNAs, is identified with a Z-score=2.33. **(B)** Proteins used as testing for our approach, experimentally validated to interact with DENV sfRNA (*Michalski et al.)* [21] and 3’ UTR (*Liao et al.*) [22]). **(C)** Barplot showing how the experimentally validated proteins interacting with DENV sfRNA are predicted by catRAPID. For each protein, we selected the best 10 interactions (Z-score) against all DENV sfRNAs. We then checked how these interactions fall inside the distribution of the human proteome interacting with DENV fragments. The proteins are well-predicted, with ∼70% of the interactions falling in the top 10% of all the ranked interactions between DENV sfRNAs and the human proteome. **(D)** Barplot showing how the experimentally validated proteins interacting with DENV 3’ UTR are predicted by catRAPID. For each protein, we selected the best 10 interactions (Z-score) against all DENV sfRNAs. We then checked how these interactions fall inside the distribution of the human proteome interacting with DENV fragments. The proteins are well-predicted, with ∼80% of the interactions falling in the top 5% of all the ranked interactions between DENV sfRNAs and the human proteome.

### GO enrichment analysis

We used GOrilla for the main GO enrichment analysis, using the entire human proteome as background [29]. The obtained p-values of the selected GOs were used as main input to draw figures.

### eCLIP analysis and the BPI index

RNA interactions for 151 RBPs were retrieved from eCLIP experiments performed in K562 and HepG2 cell lines. In order to measure the fraction of protein binders for each transcript, we applied stringent cut-offs [−log10(*p*-value) > 5 and log2(fold_enrichment) > 3] as suggested in the original paper [30]. The coordinates of the peaks were mapped to human transcripts using the GRCh38 reference genome. From these interactions, we retrieved the list of the 100 most contacted transcripts.

We implemented a Binding Promiscuity Index (BPI) to understand if a RNA molecule has a really high-number of interactions in our dataset. The BPI is calculated as the number of strong interactions (Z-score>1.5) in our dataset normalised by the number of transcripts present in each viral species:

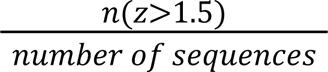

where *z* is the Z-score and *n* is the number of times this score is >1.5. We did not normalize for the sequence length since every sequence in our dataset is of 500 nucleotides. As control, we selected the last 500 nt at the 3’ UTR of the most interacting 100 mRNAs, plus, lncRNAs and mRNAs downloaded from BioMart (Ensembl version 111) of exactly 500 nt in length.

### Searching for experimental-based RBP motives

We collected conserved RNA binding motives coming from different experimental techniques (eCLIP, Bind-n-Seq, PDB, etc.) [31]. These 5-nucleotide long motives were obtained from human RBPs or, when not available, orthologous RBPs with which they share at least 70% sequence identity. Each motif is represented by a position-specific probability matrix in MEME format, for a total of 73 motives.

We used *Fimo* web server [32] to search for the previously collected motives inside our sfRNAs. *Fimo* assigned to each motive a P-value and a score. The higher the score, the higher the confidence of a motif to be inside the sequence. In our analysis, we used increasing P-value thresholds to define high-confidence motives (P<0.001 and P<0.01) and then ranked the resulting occurrences by the *Fimo* score. Depending on the threshold applied, we were able to identify a different number of highly interacting DENV proteins (5 and 14 respectively; on a total of 15 proteins with a known experimental motif).

### Granule forming predictions

We used the 97 WNV-specific protein sequences as input for *catGRANULE*, an algorithm to predict the granule-forming propensity of selected proteins [33]. Proteins with a score>0 have a propensity to be involved in granule formation. To study the significance of these findings, we then selected from the RBPome 97 random proteins, and checked how many of them have a score>0. We did this random sampling for 10,000 times and used that information to build the p-value for the 97 WNV-specific proteins.

### Checking the predictive power of custom protein libraries

We employed *cat*RAPID *library* [27] to build a custom library comprising only the 21 proteins highly interacting with all the flaviviruses. This library was fed to catRAPID Omics to predict the personalized interactome. We then used this library (ID: 792654) to identify potential sfRNA-forming regions in other flaviviruses. To do that, we divided the Murray Valley genome in non-overlapping regions of 500 nucleotides (KF751870; NCBI). Then, we checked how many of the 21 proteins have a Z-score > 1.5 for every region. The higher the proteins with a Z-score > 1.5, the higher the possibility of that viral region to be involved in sfRNA formation. Interested users can run the library at the following site by using the ID 792654 under custom dataset: http://service.tartaglialab.com/update_submission/806477/40da01a38d

## Results

### Selection of representative sfRNAs for five flaviviruses

To understand mechanisms associated with flavivirus infection, we computed a large set of interactions between sfRNAs and human RNA-binding proteins (RBPs). We first downloaded >8000 genomes of the best known flaviviruses (DENV, ZIKV, WNV, JEV, YFV; according to CDC: www.cdc.gov/vhf/virus-families/flaviviridae.html) available from different sources (**Table 1**). To select viral fragments at the 3’ UTR encoding for the sfRNAs, we used information coming from the stalling region of XRN1. We note that the complex secondary structure of ∼70 nucleotides responsible for blocking XRN1 cannot be converted into motives (scannable on new sequences) by *RNAinverse* due to its complexity [34]. For this reason, we used the known coordinates of where this complex secondary structure is located according to DENV and WNV literature [16], and used them to select the fragments at the 3’ UTR of multiple DENV and WNV genomes (**Supplementary Figure 1**). While for DENV the fragments have a length of roughly 400 nt, in the case of WNV we observed a prevalence of ∼500 nt fragments. Knowing that sfRNAs tend to be between 300-500 nt [16, 17], and to facilitate comparisons during the computational analysis, we used fragments of 500 nt at the 3’ UTR as representatives for sfRNAs. After selecting the 500 nt fragments, we reduced the intra-species redundancy with *CD-HIT* (90%, [23]), and used the retrieved fragments as representative sfRNAs for the following analysis on DENV, ZIKV, JEV, WNV, YFV.

### General characteristics of the sfRNAs

After removing sequence redundancy from our dataset, the average pairwise sequence identity among all flavivirus genomes is approximately 40% (see **Supplementary Figure 2A**). The multiple alignment analysis reveals that the regions of similarity are predominantly located at the 5’ ends of the sfRNAs, whereas the 3’ ends are more divergent (see **Supplementary Figure 2B**). These findings imply that the shared mechanisms of sfRNAs across different viruses are not solely determined by their sequence similarity, particularly after selecting representative fragments with reduced sequence similarity. Notably, the 3’ UTR region, which encodes for sfRNA formation, has been shown to be stable in DENV, both through predictions and experimental validations[35]. By focusing specifically on the 500 nt fragments in the five different flaviviruses, we find them to be highly structured based on the predicted secondary structure consensus profile, as expected from literature (**Supplementary Figure 2C**, [16, 26]). Since the secondary structure is a key element for the stalling of XRN1, complex secondary structures are directly linked to the sfRNAs activity.

### TRIM25 and other known proteins binding sfRNAs

We utilized the *cat*RAPID *omics* [27] algorithm to construct an *in silico* interactome of sfRNAs derived from more than 8,000 flavivirus genomes. This analysis generated approximately 350,000 interactions with human RNA-binding proteins (RBPs), of which around 200,000 interactions were specifically associated with the Dengue virus (DENV) (**Figure 1A**). Strong protein-RNA interactions are characterized by a Z-score > 1.5, in agreement with previous studies [28, 36]. To verify the accuracy of our predictions, we referenced known human proteins that interact with DENV sfRNAs. An example of such a protein is TRIM25, which is known to bind to DENV sfRNA [37]. This binding has an anti host-defense function for DENV by inhibiting interferon expression. We predict TRIM25 as a high-level interactor of DENV sfRNA, with a Z-score of 2.33, falling in the top 5% of all the ranked interactions between DENV sfRNAs and human proteins (**Figure 1A**).

To further validate our predictions, we used a set of experimental-validated proteins from previous studies, including proteins interacting with the sfRNA and the 3’ UTR of DENV [21, 22]. We selected the proteins specifically interacting with DENV sfRNA (21 proteins; [21]) and the high-specific proteins interacting with the DENV 3’ UTR (27 proteins; Material and Methods: Selecting and comparing experimental proteins; [22]). We note that the two sets of highly-specific proteins have only one protein in common (**Figure 1B**). After ranking all the predicted DENV interactions of our dataset, we checked for each experimental-validated protein how the best 10 interactions against DENV sfRNAs fall inside the complete distribution of human proteins interacting with DENV fragments (**Figure 1C, D**). The experimentally validated proteins interacting with DENV sfRNA are very well identified by our method, with 70% of the interactions falling in the top 10% ranked interactions and 60% of them are also in the top 5% (**Figure 1C**). The predictions are even more significant for the experimentally validated proteins interacting with DENV 3’ UTR, with 80% of the predicted interactions falling in the top 5%, and around 50% of them also falling in the top 1% of all the interactions between DENV fragments and the entire human proteome (**Figure 1D**). We further expanded our analysis by comparing our highly-interacting DENV proteins (Z-score > 1.5) with known binding motives collected from different experimental techniques (eCLIP, Bind-n-Seq, etc…; **Supplementary Figure 3**; [31]). Interestingly, we identified 15 proteins from our set with also an experimentally validated motif (5 nt motives). Moreover, if we check the presence of the motives on our set of sfRNAs, 5/15 proteins have a motif identified on the sfRNAs (p-value<0.001), and 14/15 with a less stringent p-value (p<0.01). Interestingly, if we rank the motive-identified protein by *Fimo* score (tool employed to identify sequence-based motives, [32]), the first protein identified is DHX58, a helicase mediating the antiviral signaling [38].

These results highlight the power of our predictions: we are not only able to correctly identify the binding of TRIM25 with DENV sfRNA, but our results also correctly classify two slightly-overlapping sets of experimental-validated proteins coming from two different studies. Moreover, we were able to provide a huge amount of novel high-confidence interactions, highlighting the potential of our analysis to further characterize sfRNAs.

### Expanding the sfRNA interactome with human proteins

To further expand the list of human proteins interacting with sfRNAs, and to consolidate the role of sfRNAs as anti host-defense mechanism, we studied the complete *in silico* interactome of the sfRNA of DENV, ZIKV, JEV, YFV, and WNV. In this analysis, we focused on proteins with a high interaction propensity, selecting only those with a Z-score greater than 1.5 (**Supplementary Table 1**), This selective approach allowed us to pinpoint proteins that are specific to each virus species as well as core proteins that are common across multiple flaviviruses, thereby providing insights into both unique and shared interaction patterns (**Figure 2A**). DENV has >200 highly-interacting proteins, with 47 proteins that are specific only to DENV. This is in contrast with ZIKV, which only has 34 highly-interacting proteins and zero specific proteins. WNV shows the highest number of highly-interacting proteins (>400), with 97 proteins specific only to WNV. By checking proteins highly-interacting with sfRNAs of all the flaviviruses, we identify five proteins (DDX1, NKRF, CSTF3, TRM1L, NUFP2; **Figure 2B**). Among these five proteins, we identify DDX1, an important helicase involved in host-defense mechanisms during viral infections [39]. NKRF is a regulator of DHX15, another RNA helicase involved in RNA processing and antiviral innate immunity [40], which was also seen to be inhibited by miR-301a during JEV infection [41]. Another example is NUFP2, a FMR1-interacting protein directly antagonized by the sfRNA of ZIKV [42].

**Figure 2:**
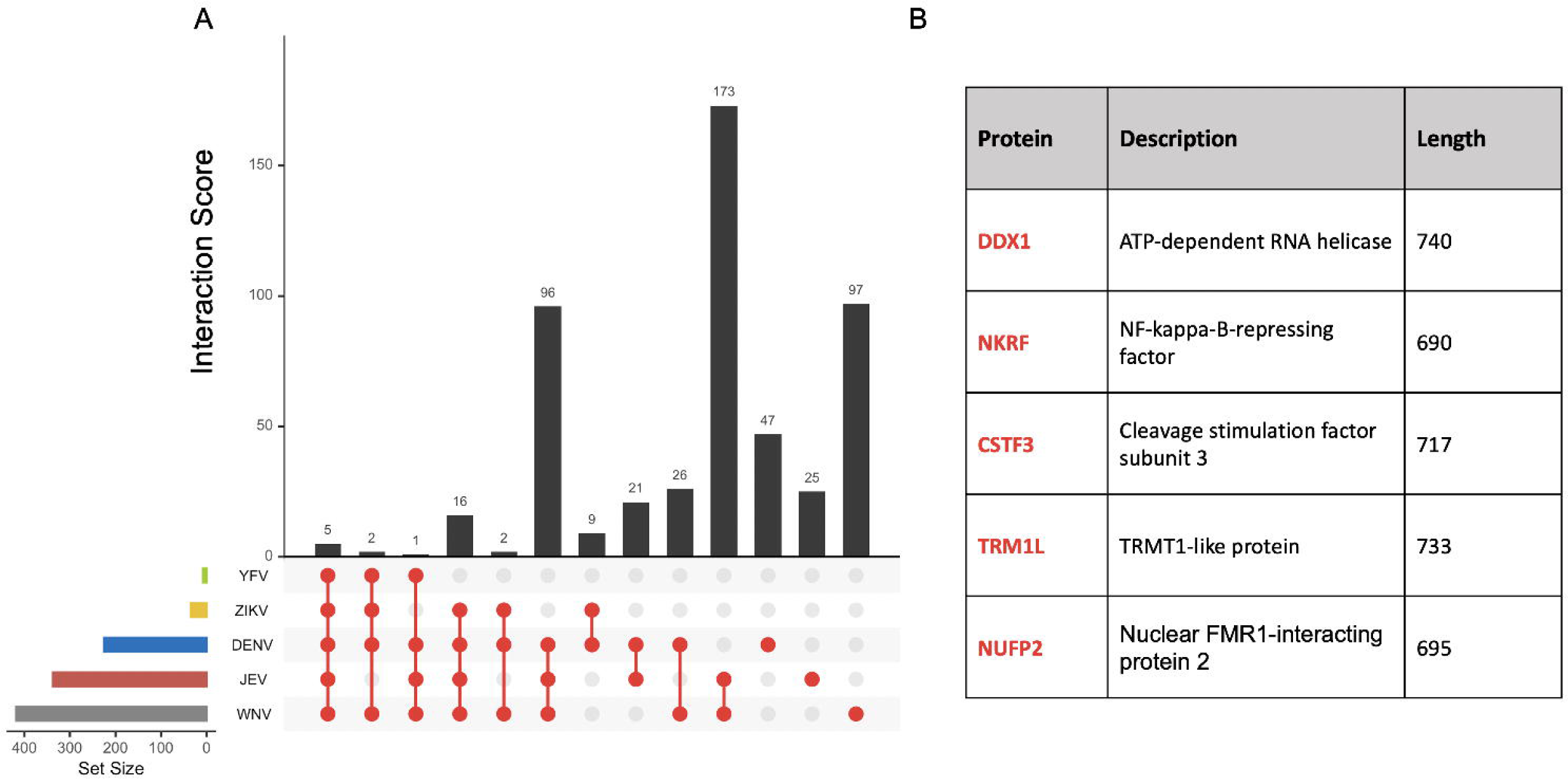
**(A)** Upset plot showing the highly interacting proteins (Z-score > 1.5) in common between the 5 different flaviviruses. The bottom-left barplot shows the total high-interacting proteins for each virus. The red dots highlight the intersection reported in the upper barplot. Five proteins are highly interacting with all the sfRNAs. **(B)** Table highlighting the characteristics of the five proteins in common between the five different flaviviruses.

To further validate the specificity of these proteins for the 3’ region of flaviviruses, we checked the interactions between these 5 proteins and the 3’ region (500 nt) of HIV-1 (GenBank: AF033819.3). None of the sfRNA-specific proteins show Z-score > 1.5 in HIV. The fact that these five proteins are found highly interacting with the sfRNAs of all the flaviviruses may indicate that the presence of sfRNAs could be a common mechanism employed by these viruses to bypass the host immune defenses, either by directly binding to important factors such as RNA helicases or by hijacking regulators of those proteins.

To expand our selection of core sfRNA-interacting proteins, since YFV has only eight highly interactive proteins, we decided to restrict the analysis by selecting the proteins in common between the other four flaviviruses. For this reason, we focused on the 21 proteins highly interacting with DENV, ZIKV, JEV, WNV (**Figure 3A; Supplementary Table 2**). Also in this case, none of the 21 proteins was found highly-interacting in HIV-1, highlighting the specificity of these proteins. By looking at the biological processes in which these proteins are involved, we found a significant enrichment for RNA splicing, processing, metabolism, and ribonucleoprotein complex assembly (**Figure 3B, C**). This enrichment already highlights the importance of sfRNAs for anti host-defense mechanisms, considering how altering these cellular host processes could disrupt the cellular machinery, promoting viral fitness. RNA splicing was already identified as the mechanism comprising the largest group of human interacting proteins, hence appearing as a highly disrupted mechanism by sfRNA presence [21]. Overall, we found evidence in literature related to flaviviruses and potential anti host-defense functions for the majority of the proteins in this set. For example, HMGN2, CSTF, and RMB39 are differently regulated upon flavivirus infection, especially DENV [21, 43, 44]. We also identified several proteins involved in splicing, including LSM2 and CCNL2. Surprisingly, we also identified SRP54 and SRP9, pro-viral proteins and negative regulators of the IFN response in this set [45]. Experiments are needed to understand the extent of these proteins binding to the sfRNAs, since their presence is supporting viral fitness.

**Figure 3:**
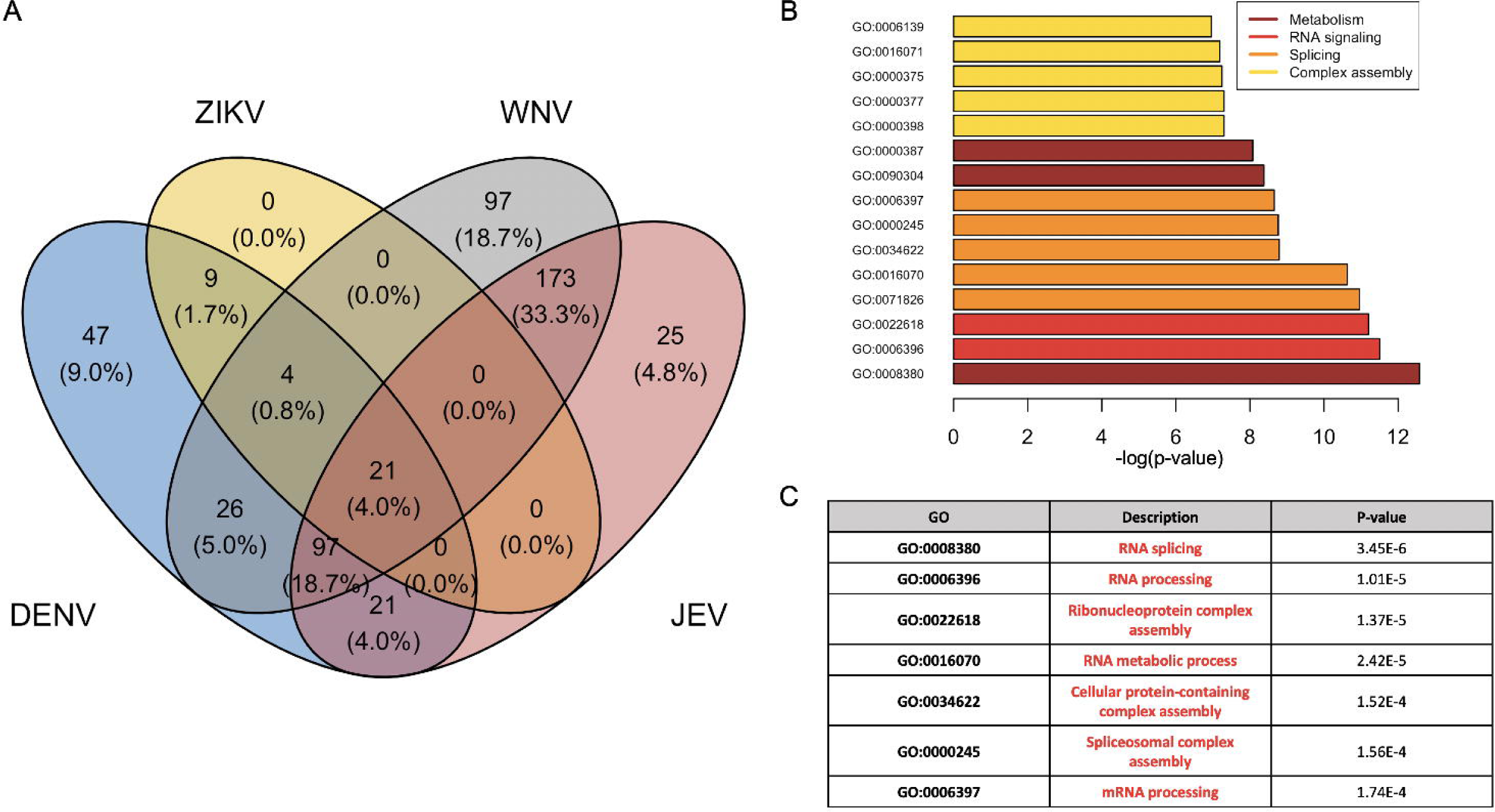
**(A)** Venn diagram showing the high-interacting proteins (Z-score > 1.5) in common among the 4 different flaviviruses. 21 proteins are considered highly interactive with all the viruses. The percentage represents the reported fraction against the total amount of proteins. **(B)** Barplot showing the p-value of the most significant GOs identified from the enrichment of the 21 common proteins. Terms associated with similar mechanisms share the same colours. **(C)** Table extracted from GOrilla showing specific significant GO terms from the enrichment of the 21 common proteins.

The protein L10K, produced by the gene C19orf53, is an interferon-stimulated gene (ISG) product still not well characterized. However, C19orf53 is found in IFN cDNA libraries together with C19orf66, a recently characterized ISG involved in the antiviral response against DENV and JEV [46, 47]. Because of these similarities, L10K is a very promising candidate for further studies to better understand the sfRNAs contribution and the interferon response of the flaviviruses.

### Couple- and species-specific proteins and disrupted mechanisms

JEV and WNV (JEV-WNV) share the highest number of common highly-interacting proteins (173 proteins), while DENV-JEV and DENV-WNV have a similar number of common highly-interactive proteins (21 and 26 respectively; **Figure 4**). As in the previous analysis, we were able to identify known disrupted mechanisms, including proteins associated with RNA processing and RNA metabolism **(Supplementary Figure 4)**. With the exception of the couple DENV-WNV (**Figure 4A**), by looking at the molecular function of the couple-specific proteins, we identified more specific processes. The G protein-coupled receptor signaling pathway is identified as enriched in the proteins specific to WNV-JEV (**Figure 4B**), a mechanism known to be hijacked during viral infection and tumorigenesis [48]. Moreover, proteins involved in chemokines activity and signaling are also highly binding WNV-JEV sfRNA, a very interesting result since chemokines are crucial for the control of viral infections and part of the IFN cascade [49]. RNA helicases are highly binding the sfRNAs of DENV-JEV, a class of crucial molecules for their antiviral activity, already reported as a disrupted mechanism for the core 21 proteins (**Figure 4C**). Moreover, these sfRNAs seem to compete with proteins binding the 3’ UTR of host mRNAs, thus disrupting the post-transcriptional regulation.

**Figure 4:**
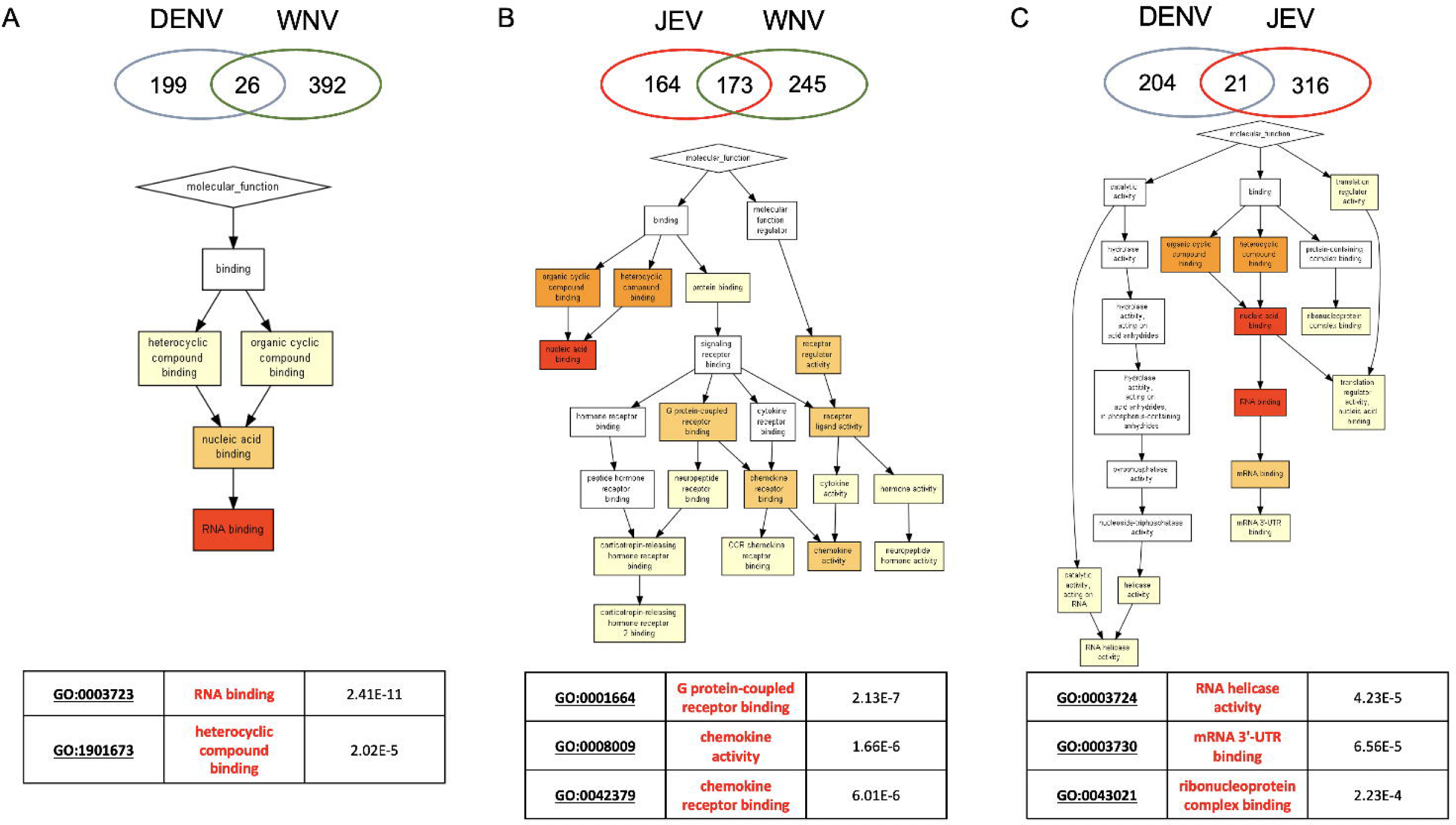
**(A)** Enrichment analysis of the 26 proteins highly interacting specifically with DENV and WNV. The GO pathway and summary table were extracted from GOrilla. **(B)** Enrichment analysis of the 173 proteins highly interacting specifically with JEV and WNV. The GO pathway and summary table were extracted from GOrilla. **(C)** Enrichment analysis of the 21 proteins highly interacting specifically with DENV and JEV. The GO pathway and summary table were extracted from GOrilla.

We also concentrated on species-specific proteins, those that bind with high affinity exclusively to a particular virus. As previously reported in our study (see section: Expanding the sfRNA interactome with human proteins), WNV shows the highest number of species-specific proteins (**Supplementary Figure 5**). These proteins are involved in RNA metabolism and RNA processing, but are also implicated in more specific functions, for example as structural constituents of ribosomes (**Supplementary Figure 6**). Interestingly, DENV-specific proteins are involved in the binding with the poly-U, poly-A, and 3’ UTR of the mRNAs (**Supplementary Figure 7**), highlighting again the possible competition with host mRNAs for the binding. Moreover, viruses have different mechanisms to disrupt the poly-A binding in order to inhibit host-translation, for example by cleaving or displacing proteins [50]. In this case, DENV could employ a displacing strategy through the sfRNA.

These results coming from couple- and species-specific proteins shed light on mechanisms such as the G protein-coupled signaling, chemokines activity and the heterocyclic compound binding, the latter being particularly relevant considering that some of these molecules have been discovered experimentally to inhibit DENV infection in cell culture [51]. These mechanisms complement the list of processes discovered in the previous analysis of the core-proteins, common to all flaviviruses. This shows how crucial mechanisms such as RNA processing and metabolism can be altered during viral infections. Altogether, our results highlight the importance of the sfRNAs to disrupt crucial cellular mechanisms, inhibiting the host-defense system and promoting viral fitness and translation. The results of our enrichment analysis coming from >8000 flaviviral genomes comprising 5 different viruses are summarized in **Table 2**.

### Binding promiscuity index and sfRNAs as *protein sponges*

sfRNAs compromise the host-defense immunity, not only by disrupting important cellular mechanisms but also by directly altering the IFN-mediated immune response and other antiviral-related processes. We checked whether this hijacking activity could be caused by the sfRNAs acting as *protein sponges*. For this to be the case, sfRNAs should have a high-number of promiscuous but stable interactions. To validate this, first, we built a binding promiscuity index (BPI) for each flavivirus as the number of predicted strong interactions (Z-score > 1.5) normalized for the number of sequences in each set. We did not normalize for the RNA length since all the fragments were of 500 nt. While DENV and ZIKV show a low BPI, JEV and especially WNV have a high BPI. In order to establish the power of the BPI index to show that JEV and WNV sfRNAs have the potential to bind many host proteins, we employed positive controls based on available experimental data. We analyzed the RNA molecules collected from eCLIP experiments (**Figure 5A**, **Materials and Methods**; [52]), ranking these RNAs for the number of significant protein contacts. While the majority of the RNAs have very few contacts (**Figure 5B**), some molecules show a high number of protein interactions. To test our approach, we selected the 3’ region (500 nt) of the 100 mRNAs with the highest interaction with proteins, according to eCLIP data (**Figure 5B**) and used these fragments to compute RBP interactions with *ca*tRAPID *omics* [27], applying the same procedure of the sfRNAs. Then, we used the same approach to compute the BPI for these RNAs. We further validated the BPI by selecting lncRNAs and coding RNAs exactly 500-nt long. While the most interacting RNAs show the highest BPI, as expected, WNV has a higher BPI than lncRNAs and mRNAs of the same length (**Figure 5C).** These results suggest how sfRNAs, especially WNV, tend to have a high number of protein interactions. Moreover, the sfRNAs compete for the binding with the mRNAs, showing a higher BPI than mRNAs of the same length, highlighting even more their anti host-defense functions.

**Figure 5:**
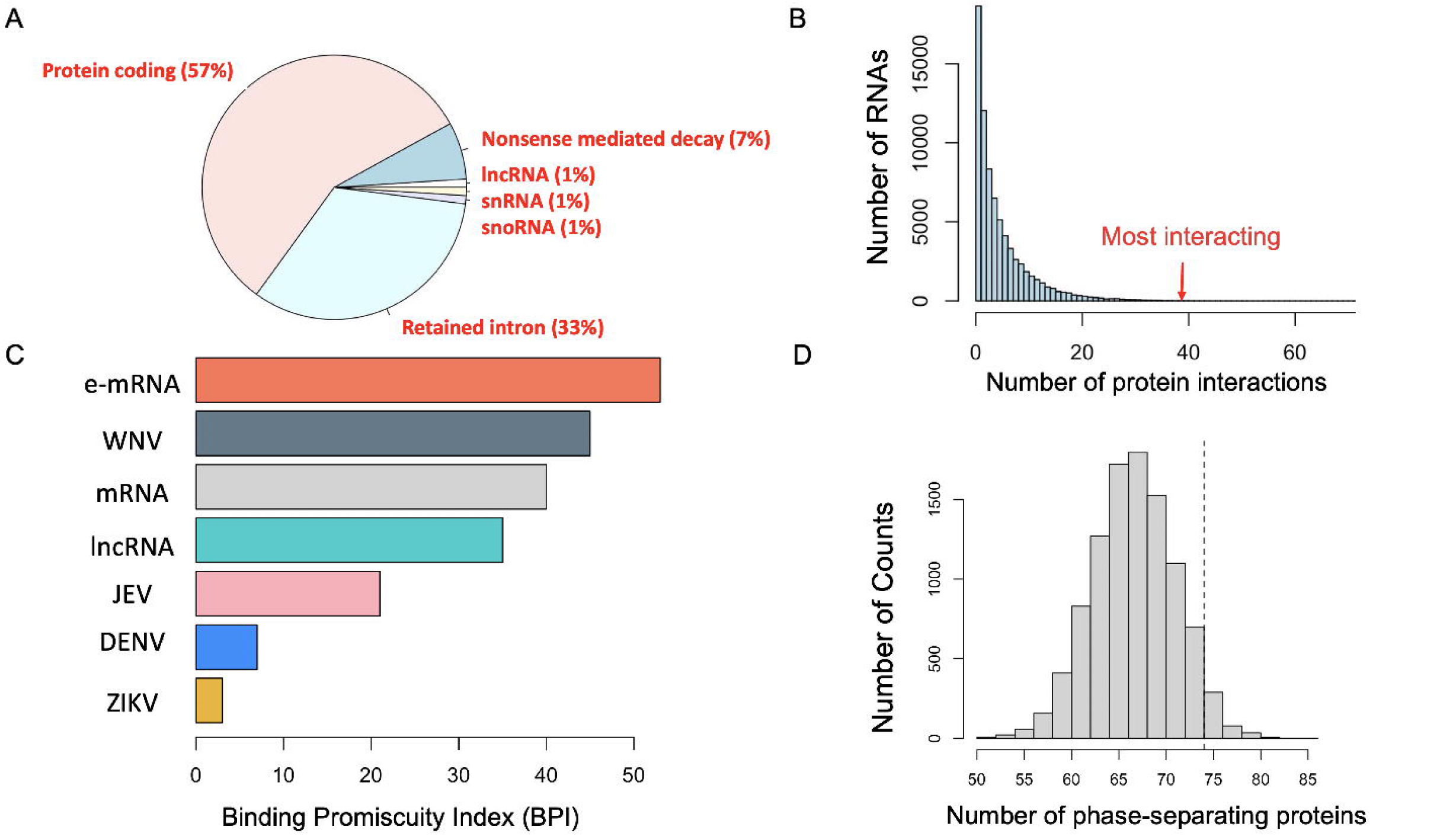
**(A)** Pie Chart of the eCLIP data, showing the biotype classes of the 100 RNAs mostly contacted by proteins. **(B)** Histogram of the eCLIP data, showing the number of protein interactions with RNAs. The majority of the RNAs tend to have very few interactions, according to eCLIP data. The average of the 100 most interacting mRNAs is highlighted in the plot. **(C)** Horizontal bar plot showing the Binding Promiscuity Index (BPI) computed for the 5 flaviviruses. The 3’ end of the 100 mRNAs mostly interacting with proteins (e-mRNAs), according to eCLIP data, plus lncRNAs and mRNAs of 500 nucleotides are used as control. **(D)** *cat*GRANULE significance analysis. For 10,000 times, 97 random proteins were selected from the RBPome, and we checked how many of them have a score>0, according to *cat*GRANULE. The dashed line represents the threshold of the WNV-specific proteins having a score>0.

One of the cellular mechanisms to fight viral infections is the formation of stress granules, which occurs when the viral RNA is sensed by specific proteins such as RIG-I [38], subsequently stalling the rate of mRNAs translation. Unlike solid-like aggregates [53, 54], stress granules rapidly assemble to protect the cell during infection and dissolve quickly afterward [28]. In this context, to investigate the anti innate immunity potential of WNV sfRNA, we studied the granule-forming propensity of the >90 proteins highly-interacting specifically with WNV. This is relevant because, for example, the antiviral mechanism of stress granule formation is inhibited during WNV infection [55, 56]. We used *cat*GRANULE [33] to predict the propensity of these proteins to form granules. Interestingly, >75% of the WNV-specific proteins exhibit a propensity to undergo phase separation. (**Supplementary Figure 8**). This result is significant when compared to similar random distributions (p-value<0.05; **Figure 5D; Materials and Methods**: Granule forming predictions). Establishing strong bindings with proteins involved in stress granules or other phase-separated complexes could represent an additional WNV strategy against the host defenses, where the sfRNAs could bind and sequester important components of these organelles to avoid their formation and ensure viral fitness.

### Predicting sfRNA-forming regions employing a subset of human protein interactors

We identified a set of 21 proteins highly-specific and interacting with the sfRNAs of DENV, WNV, JEV, and ZIKV. The specificity of these proteins for the sfRNA-forming regions could be used to further study or characterize novel or less-studied viruses. To test this hypothesis, we used *cat*RAPID *library* [27] to build a custom protein dataset for the 21 proteins to be then used in *cat*RAPID *omics* to predict the interactions. Then, we selected a rare and poorly studied flavivirus, not from Asia or South America, the Murray Valley virus (MVV). MVV is an arbovirus from Australia, forming sfRNAs and exploiting mosquito as a vector a[2, 57]. We studied the complete set of interactions between MVV and the 21 proteins previously identified. For all the fragments of 500 nt, we checked how many of the 21 proteins were highly-interacting (Z-score > 1.5) in every specific region (**Figure 6**). Interestingly, the 3’ UTR regions is the only one showing high-interactions with all the 21 proteins, highlighting how this set of proteins can in fact identify sfRNA-forming regions. Moreover, if we perform the same analysis on a non-flavivirus, in this case HIV-1 divided in fragments of 500 nucleotides, we identified only 2/21 highly-interacting (Z-score > 1.5) proteins in the region region with the highest number of protein-interactions, and not located in the 3’ UTR (**Supplementary Figure 9**). This result shows how this set of 21 proteins can be used to further study RNA viruses in order to identify regions encoding for sfRNA formation. We propose that this set of proteins could be used to identify other anti host-defense regions in the genome of RNA viruses. For this reason, users can select the custom-library ID (792654) to run specific *catRAPID* analysis on these proteins (see **Material and Methods**: Running the predictive libraries).

**Figure 6:**
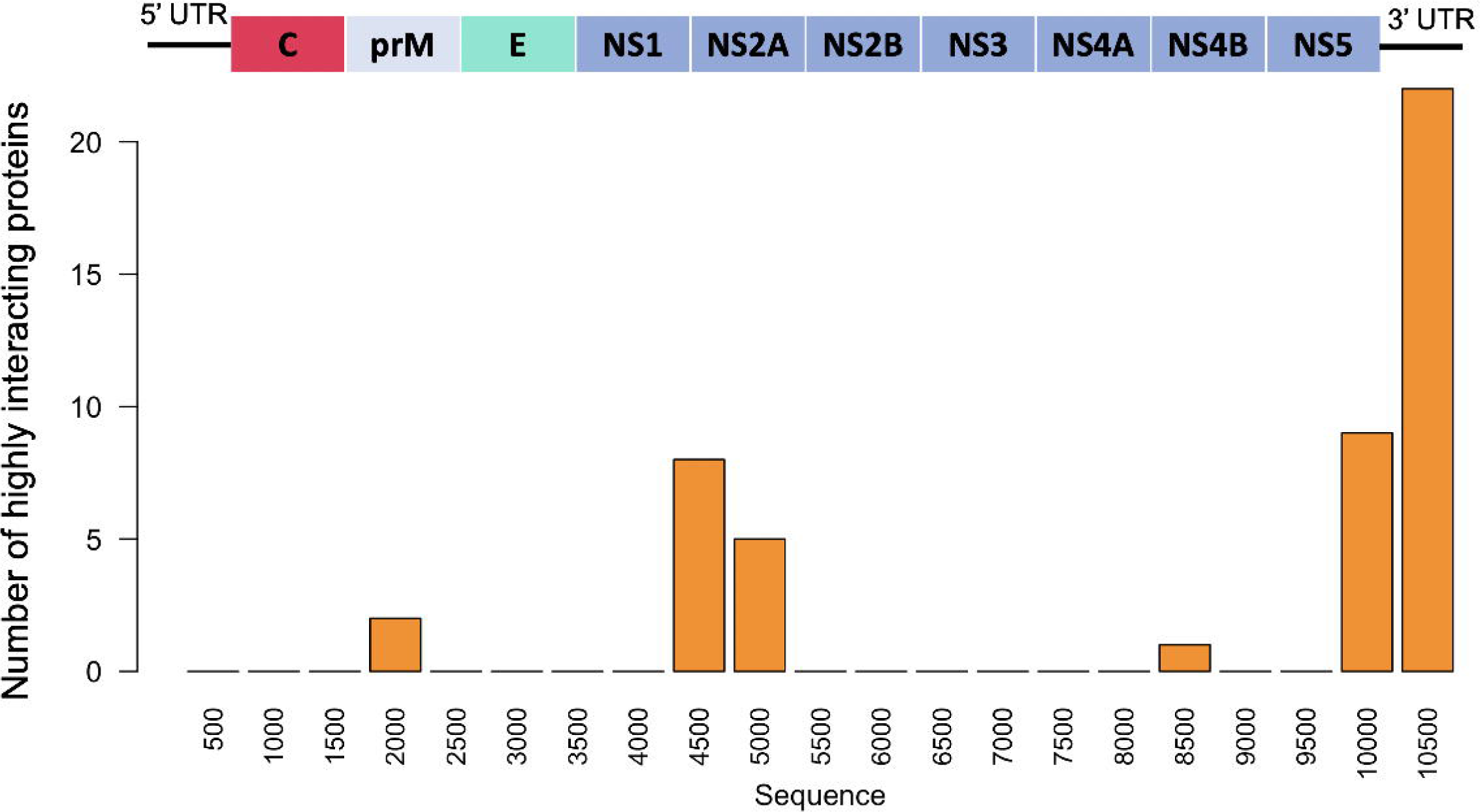
The genome of the Murray Valley virus (MVV) was divided in fragments of 500 nucleotides. For each region, we highlighted the number of highly interacting proteins (Z-score > 1.5) from the pool of the 21 sfRNA-specific proteins. The 3’ region is the one highly interacting with all the proteins.

## Discussion

Viruses employ various mechanisms to evade the host’s innate immune response, and flaviviruses are no exception, utilizing several strategies to disrupt cellular functions or circumvent the IFN-mediated response. A characteristic mechanism of flaviviruses involves the formation of sfRNAs, which result from the stalling of the XRN1 enzyme at the 3’ UTR of flaviviral genomes. These RNA fragments have been shown to play a crucial role in countering host-defense mechanisms. However, the full extent of sfRNAs’ roles was not completely understood due to the limited number of flaviviral genomes analyzed and the few common candidate proteins identified in different studies.

In our research, we analyzed over 8,000 genomes from five different flaviviruses (DENV, ZIKV, WNV, JEV, YFV) to examine the predicted in-silico interactome with human proteins. We approximated sfRNAs using 500-nucleotide fragments based on the distribution of fragments generated from XRN1-stalling coordinates. Our in-silico interactome comprised more than 300,000 interactions between sfRNAs and human proteins. Among the most interacting proteins, we identified known sfRNA interactors, including TRIM25. To further validate our predictions, we utilized two sets of experimentally validated proteins that interact with the DENV sfRNA and its 3’ UTR [21, 22].

Supporting the high quality of our predictions, we identified that approximately 70% of the experimentally validated sfRNA-specific interactions fell within the top 10% of ranked interactions. Additionally, about 80% of the predicted interactions involving 3’-specific proteins ranked within the top 5% of all interactions between DENV fragments and the entire human RNA-binding proteome (RBPome; **Figure 1**).

We identified the interactions of each virus, including species-specific proteins: 47 proteins highly interacting specifically only with DENV, 97 with WNV, and 25 with JEV. WNV and JEV showed the highest number of coupled-specific proteins, with 173 proteins interacting only with WNV and JEV. Importantly, 21 proteins were classified as highly interacting with all the flaviviruses, excluding YFV. Among these proteins, we identified different helicases and their interactors, including DDX1, DHX58, and a regulator of DDX15. Knowing the innate antiviral activity of the RNA helicases, it is easy to speculate how the binding with the sfRNAs disrupt this host-defense mechanism, promoting viral fitness.

We computationally investigated the general mechanisms and functions of human proteins interacting with sfRNAs. The majority of highly interactive proteins are associated with RNA splicing, signaling, and metabolic processes—crucial host cellular functions that are often disrupted by viral infections. Specifically, RNA splicing is the primary mechanism involving most of the proteins bound to DENV sfRNA, as corroborated by experimental data [21]. Species-specific and couple-specific proteins tend to be linked with more targeted anti-host defense mechanisms, such as competing for binding at the 3’ UTR with host coding RNAs or disrupting the activity of RNA helicases and chemokines. These findings shed light on the multiple layers of anti-host defense mechanisms employed by sfRNAs, which can hijack and disrupt general cellular mechanisms common to all flaviviruses, while also displaying alterations specific to each viral species.

This is possible due to the numerous strong interactions that sfRNAs can establish with human proteins. We demonstrated that WNV, in particular, has a higher number of strong interactions compared to coding and long non-coding RNAs of the same length. Additionally, WNV appears to be associated with granule-forming proteins, likely to inhibit the formation of stress granules, thereby enhancing viral fitness [28, 36]. This supports our hypothesis that sfRNAs not only disrupt and hijack various cellular mechanisms but also function as protein sponges by establishing a high number of potential bindings.

## Conclusions

In this work, we computed the largest *in silico* interactome of flaviviruses to understand how the accumulation of sfRNAs in human cells can disrupt host-defense mechanisms. Our analysis provided a way to exploit the newly-identified candidate proteins. Indeed, we demonstrated that a set of 21 proteins, which interact with sfRNAs from all the different viruses, can be used as a predictive tool to identify sfRNA-forming regions in other cases, such as the Murray Valley virus—a distinct Australian flavivirus not previously included in our analysis. Researchers can also employ this set of proteins with the *cat*RAPID *library* [27] to characterize novel or understudied flaviviruses. We plan to integrate this information into other algorithms, such as *RNAvigator* [35], to utilize specific protein interactions to identify characteristic features of RNA regions.

## Abbreviations

DENV: Dengue virus
ZIKV: Zika virus
WNV: West Nile Fever virus
JEV: Japanese Encephalitis virus
YFV: Yellow Fever virus
HCV: Hepatitis C virus
RBP: RNA binding protein
UTR: Untranslated region
GO: Gene ontology
ss-RNA: Single-stranded RNA
lncRNA: Long non-coding RNA
sfRNA: Subgenomic flaviviral RNA

## Data availability

The catRAPID interactions are available as individual links in the Methods. The most interacting proteins for all the viruses are available in the Supplementary Table 1. The complete list of the 21 proteins interacting with all viruses is available in Supplementary Table 2.

## Acknowledgement

The authors would like to thank the other members of Tartaglia’s group and Roland Huber for the useful comments.

## Contributions

RDP conceived the study. RDP and GGT designed the study. RDP and AV performed the analysis and assembled the figures. RDP, AV, and GGT wrote the manuscript. All the authors read and agreed with the content of the manuscript.

## Fundings

The research leading to this work was supported by the ERC ASTRA_855923 (G.G.T.) and EIC Pathfinder IVBM4PAP_101098989 (G.G.T.).

---

**Supplementary Figure 1:** Length of the fragments from all DENV and WNV genomes generated at the 3’ UTR by using known coordinates for the XRN1 structure.

**Supplementary Figure 2: (a)** Sequence identity of all the sfRNAs fragments after removing sequence redundancy. **(b)** Multiple-alignment obtained with MAFFT of the sfRNA fragments. **(c)** Consensus secondary structure profile obtained from the CROSS predictions of all the sfRNAs fragments. Each position is the average CROSS score for all the sfRNAs in that position. A score >0 highlights a propensity for that nucleotide to be double-stranded.

**Supplementary Figure 3: (a)** Piechart showing the experiments identifying a binding motif for the 15 proteins selected in DENV. **(b)** Ranking of Fimo’s score with a p-value<0.001 of the 5 proteins identified from our predictions and having an experimental motif.

**Supplementary Figure 4:** Enrichment analysis of the couple-specific proteins equivalent to Figure 4, here showing the expanded tables of GOs.

**Supplementary Figure 5:** Enrichment analysis of the DENV-specific proteins, with the table and pathway obtained from GOrilla.

**Supplementary Figure 6:** Enrichment analysis of the JEV-specific proteins, with the table and pathway obtained from GOrilla.

**Supplementary Figure 7:** Enrichment analysis of the WNV-specific proteins, with the table and pathway obtained from GOrilla.

**Supplementary Figure 8:** Propensity to phase-separate for the >90 proteins highly interacting only with WNV, computed using catGRANULE. A positive score highlights proteins prone to phase-separate.

**Supplementary Figure 9:** The entire genome of the HIV-1 virus, divided in fragments of 500 nucleotides. For each region, we highlighted the number of highly interacting proteins (Z-score > 1.5) from the pool of the 21 sfRNA-specific proteins.

## Notes

### Competing Interest Statement

The authors have declared no competing interest.

