## Supplementary Figures for "Subgenomic flaviviral RNAs and human proteins: *in silico* exploration of anti-host defense mechanisms"

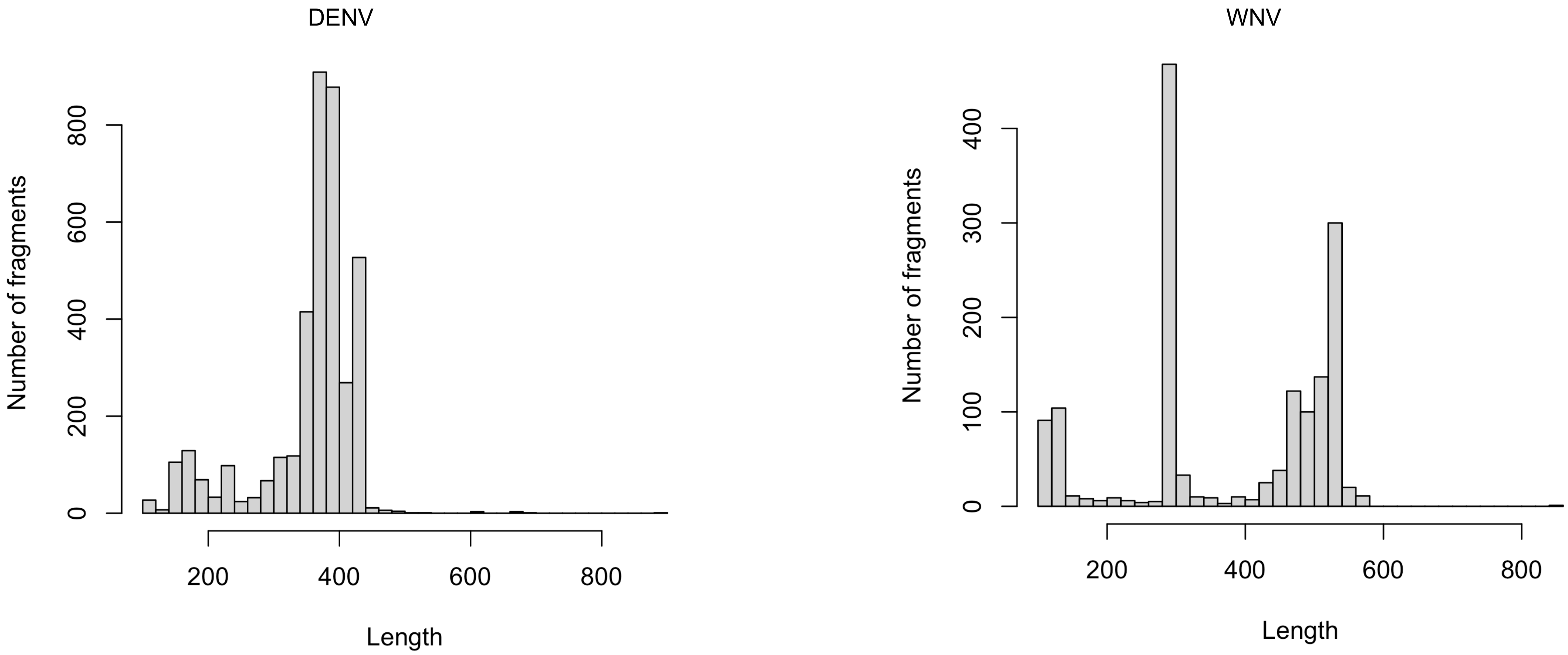
DENV
Number of fragments
0
200
400
600
800
200
400
600
800
Length
WNV
Number of fragments
0
100
200
300
400
200
400
600
800
Length

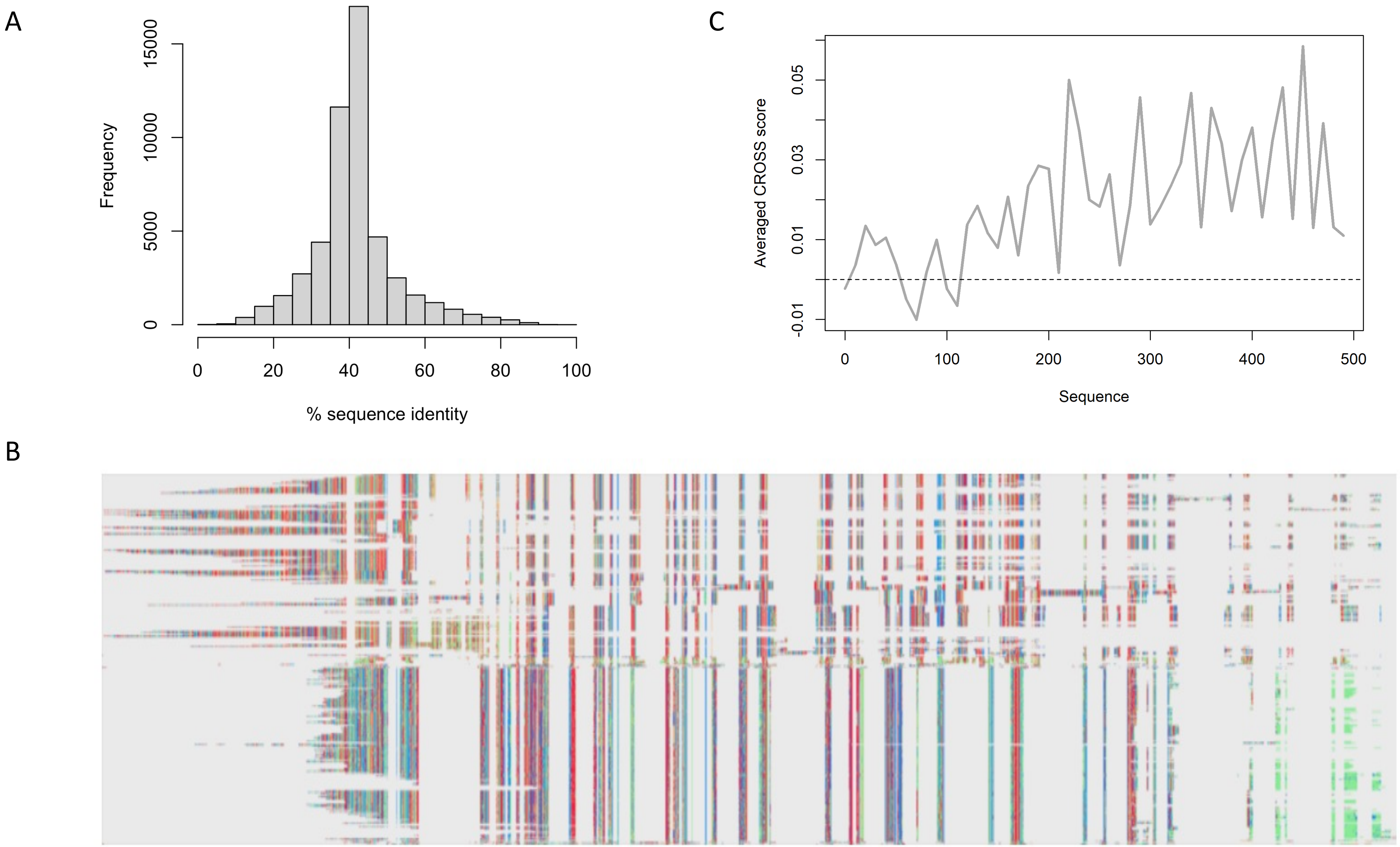
A
Frequency
0
5000
10000
15000
0
20
40
60
80
100
% sequence identity
C
Averaged CROSS score
-0.01
0.01
0.03
0.05
0
100
200
300
400
500
Sequence
B

A

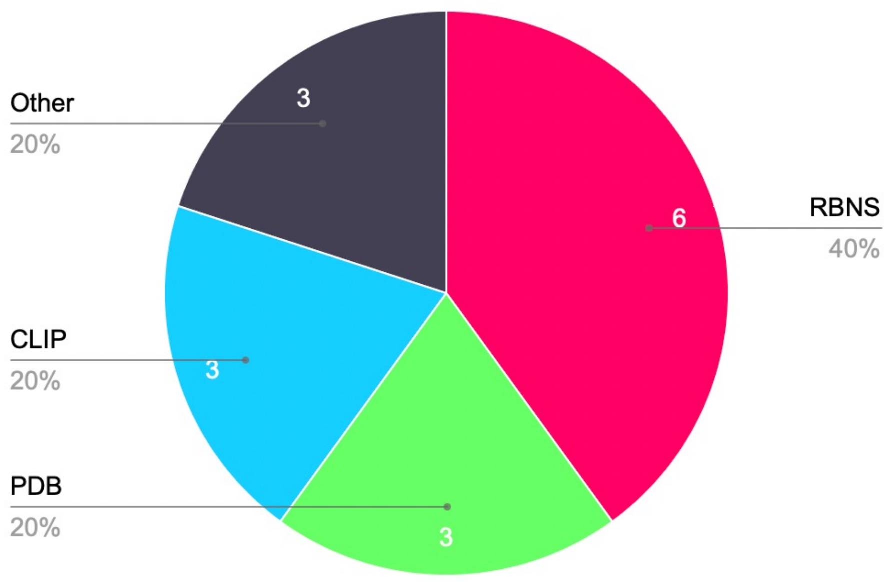

Other
3
20%
RBNS
6
40%
CLIP
3
20%
PDB
3
20%


B

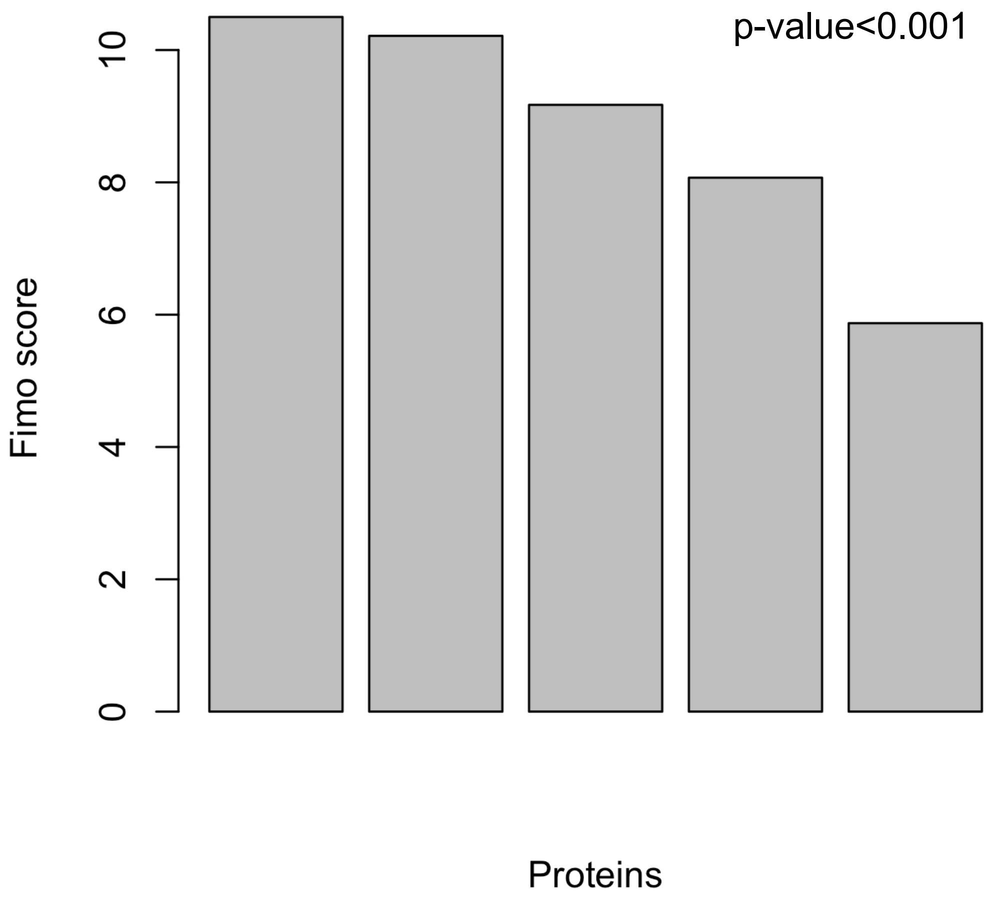

p-value<0.001
Fimo score
0
2
4
6
8
10
Proteins

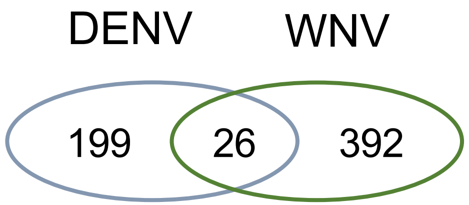


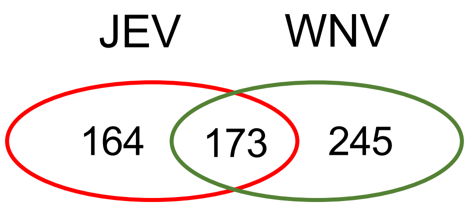


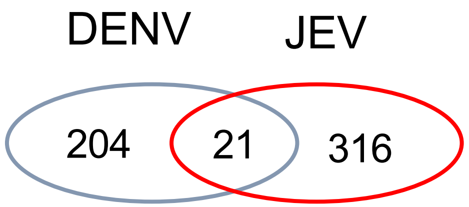


| GO term | Description | P-value |
|---|---|---|
| GO:0008380 | RNA splicing | 5.02E-8 |
| GO:0006397 | mRNA processing | 1.25E-7 |
| GO:0016071 | mRNA metabolic process | 1.21E-6 |
| GO:0006396 | RNA processing | 5.65E-6 |
| GO:0016070 | RNA metabolic process | 1.2E-5 |
| GO:0000398 | mRNA splicing, via spliceosome | 1.62E-5 |
| GO:0000377 | RNA splicing, via transesterification reactions with bulged adenosine as nucleophile | 1.62E-5 |
| GO:0000375 | RNA splicing, via transesterification reactions | 1.74E-5 |
| GO:0043484 | regulation of RNA splicing | 9.21E-5 |
| GO:0090304 | nucleic acid metabolic process | 1.18E-4 |

| GO term | Description | P-value |
|---|---|---|
| GO:0048024 | regulation of mRNA splicing, via spliceosome | 8.11E-15 |
| GO:0050684 | regulation of mRNA processing | 1.62E-14 |
| GO:0043484 | regulation of RNA splicing | 2.59E-14 |
| GO:0006396 | RNA processing | 2.11E-13 |
| GO:0006397 | mRNA processing | 2.12E-12 |
| GO:0008380 | RNA splicing | 2.51E-12 |
| GO:1903311 | regulation of mRNA metabolic process | 8.85E-11 |
| GO:0016070 | RNA metabolic process | 1.68E-10 |
| GO:0090304 | nucleic acid metabolic process | 6.1E-10 |

| GO term | Description | P-value |
|---|---|---|
| GO:0006396 | RNA processing | 4.34E-10 |
| GO:0016070 | RNA metabolic process | 1.29E-7 |
| GO:0034641 | cellular nitrogen compound metabolic process | 1.32E-7 |
| GO:0006417 | regulation of translation | 4.4E-7 |
| GO:0008380 | RNA splicing | 6.28E-7 |
| GO:0016071 | mRNA metabolic process | 7.68E-7 |
| GO:0034248 | regulation of cellular amide metabolic process | 1.26E-6 |
| GO:0006397 | mRNA processing | 1.54E-6 |
| GO:0006139 | nucleobase-containing compound metabolic process | 2.75E-6 |
| GO:0090304 | nucleic acid metabolic process | 3.38E-6 |

### DENV

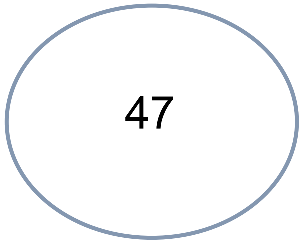


| GO term | Description | P-value |
|---|---|---|
| GO:0016070 | RNA metabolic process | 5.87E-9 |
| GO:0006396 | RNA processing | 6.48E-8 |
| GO:0090304 | nucleic acid metabolic process | 2.26E-7 |
| GO:0016071 | mRNA metabolic process | 2.72E-6 |
| GO:0009451 | RNA modification | 3.26E-6 |
| GO:0034641 | cellular nitrogen compound metabolic process | 6.67E-6 |
| GO:0006139 | nucleobase-containing compound metabolic process | 7.34E-6 |
| GO:1902373 | negative regulation of mRNA catabolic process | 1.16E-5 |

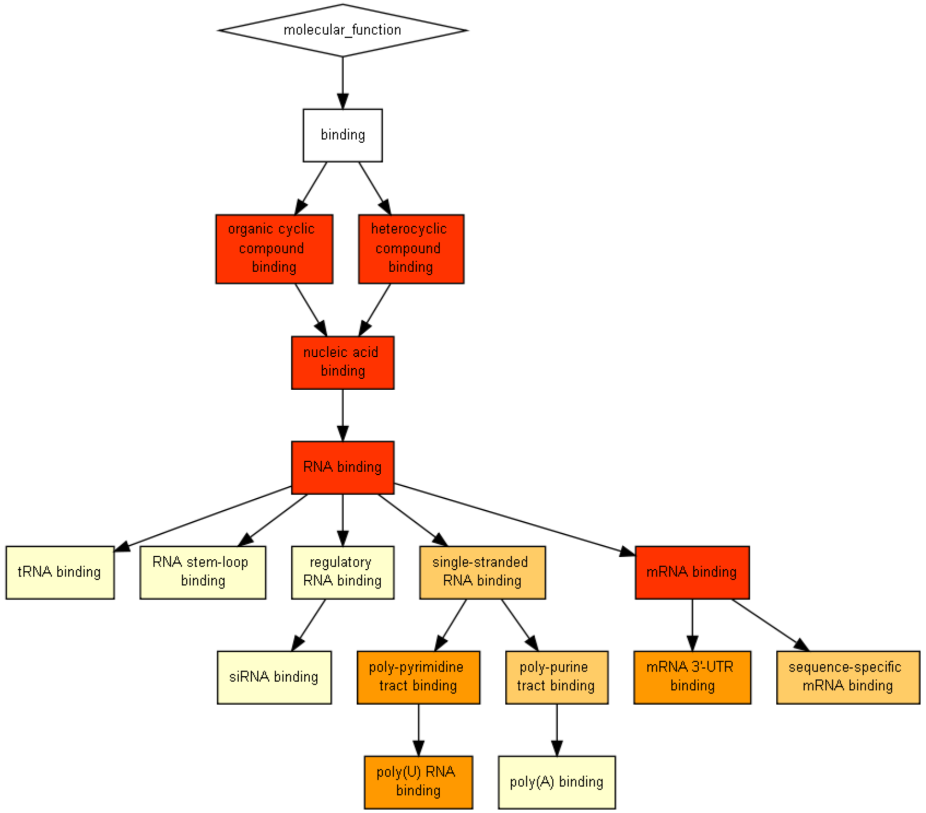

### JEV

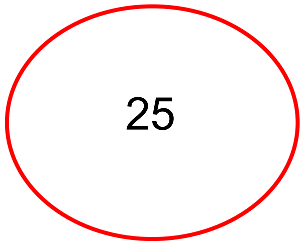


| GO term | Description | P-value |
|---|---|---|
| GO:0006406 | mRNA export from nucleus | 3.39E-4 |
| GO:0006405 | RNA export from nucleus | 6.14E-4 |
| GO:0051028 | mRNA transport | 8.24E-4 |

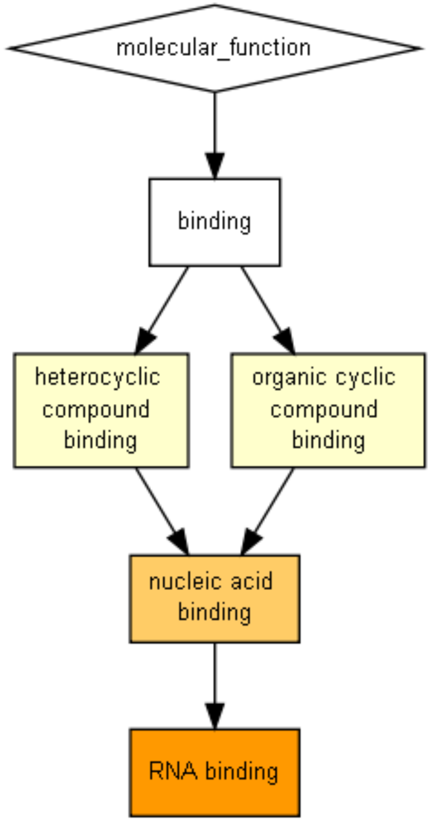

### WNV

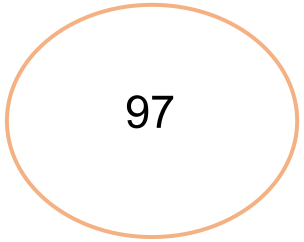


| GO term | Description | P-value |
|---|---|---|
| GO:0016071 | mRNA metabolic process | 2.96E-24 |
| GO:0006402 | mRNA catabolic process | 5E-23 |
| GO:0000956 | nuclear-transcribed mRNA catabolic process | 2.81E-22 |
| GO:0016070 | RNA metabolic process | 5.1E-22 |
| GO:0006401 | RNA catabolic process | 9.05E-22 |
| GO:0090304 | nucleic acid metabolic process | 2.41E-18 |
| GO:0010629 | negative regulation of gene expression | 5.45E-17 |
| GO:0034655 | nucleobase-containing compound catabolic process | 5.57E-17 |
| GO:0000184 | nuclear-transcribed mRNA catabolic process, nonsense-mediated decay | 1.43E-16 |
| GO:0006396 | RNA processing | 1.52E-16 |

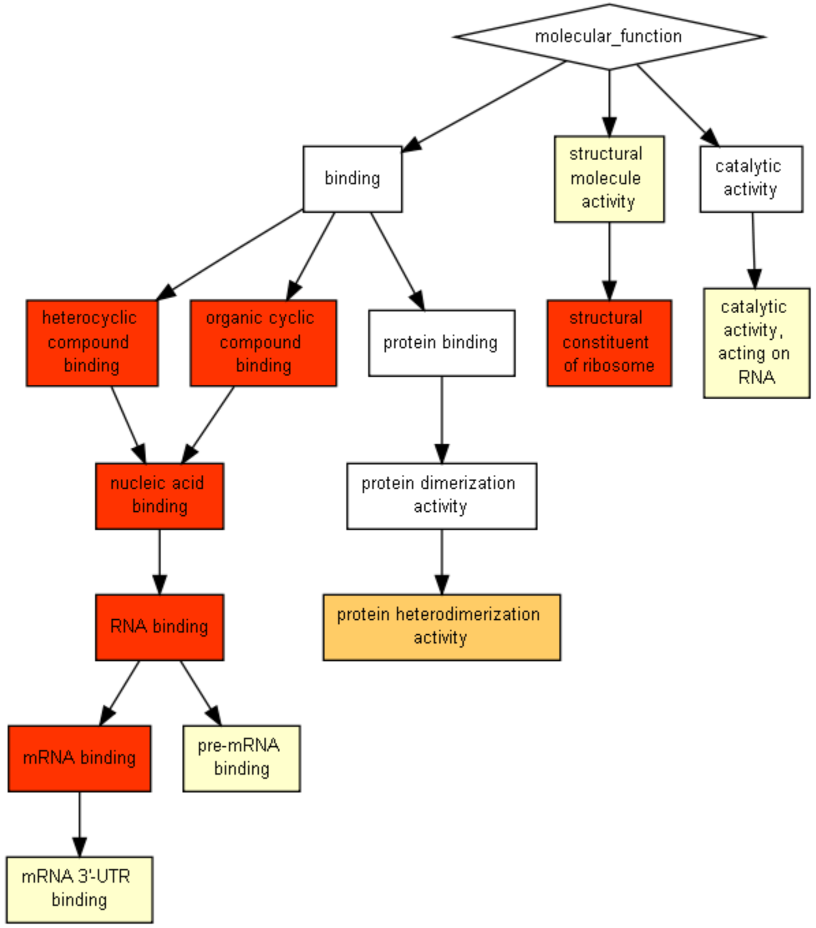

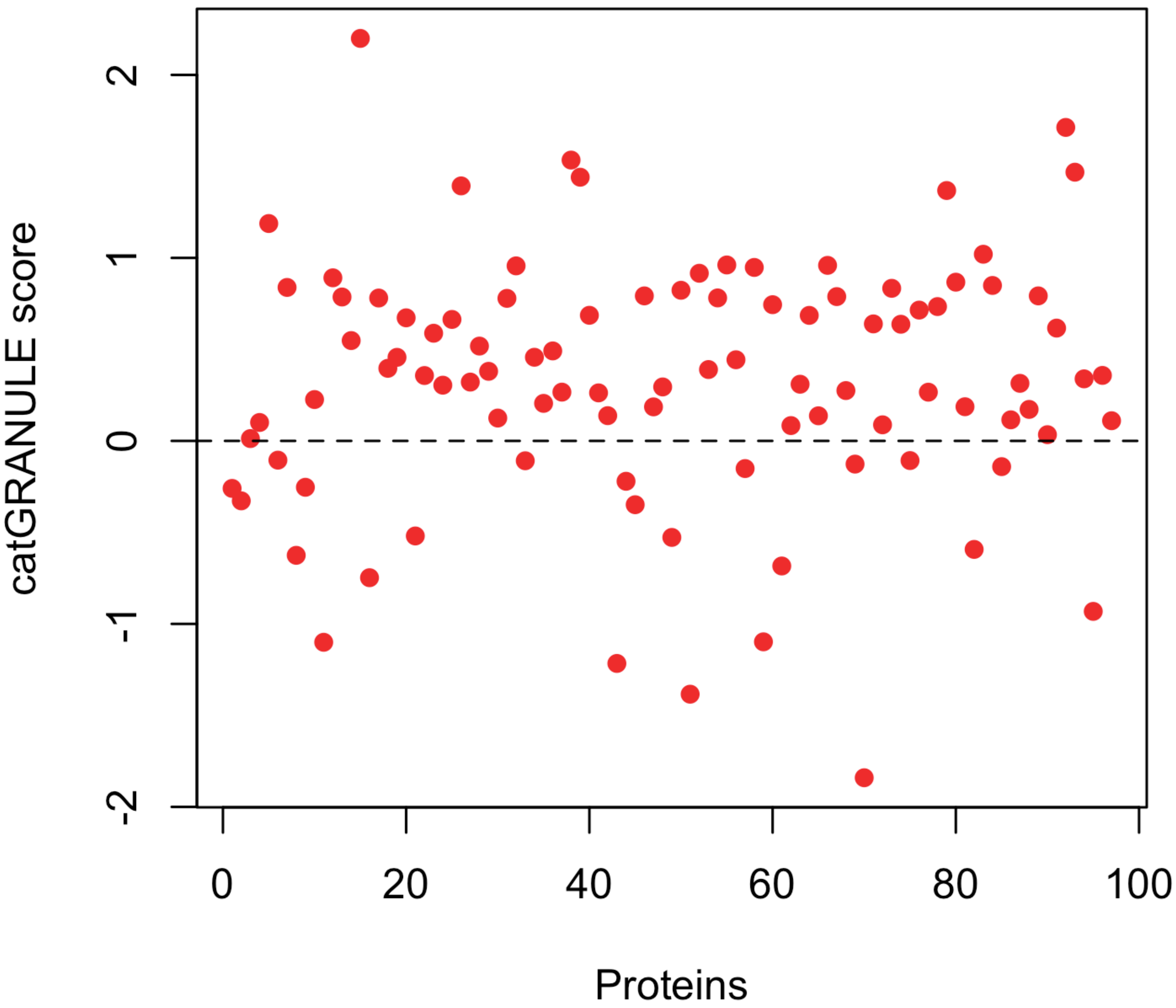

catGRANULE score
2
1
0
-1
-2
0
20
40
60
80
100
Proteins

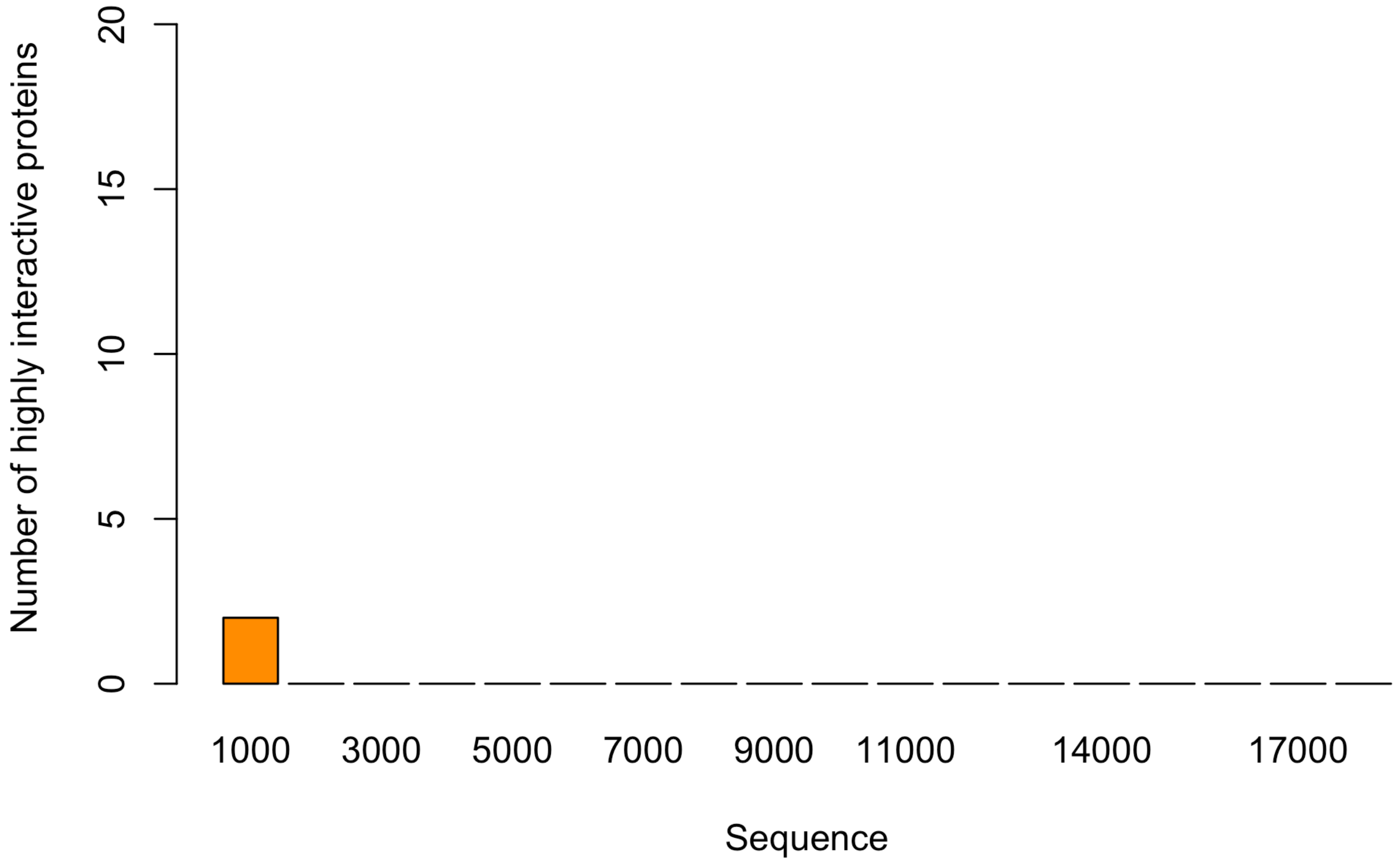

Number of highly interactive proteins
0
5
10
15
20
1000
3000
5000
7000
9000
11000
14000
17000
Sequence
